# Studies for searching factors relating rumination time around the heat season in Holstein milking cows

**DOI:** 10.1101/2022.02.25.481872

**Authors:** Takahiro Seto, Yuichi Toba

**Author notes:** Correspondence Takahiro Seto, Department of Agricultural Production, Shizuoka Professional University Junior College of Agriculture, Tomigaoka 678-1, Iwata, Shizuoka 438-8577, Japan.

## Abstract

To identify factors affecting rumination time (RT), factorial analyses in a dairy farm were performed. In univariate analyses, differences in distribution were observed between low (< 45 kg/d) and high (≥ 45 kg/d) milk yield (MY) before the heat season, early and peak lactation periods and mid and late at the start of the heat season, and 1-2 and ≥ 3 parity at the start of the heat season. Multiple regression analysis confirmed that RT was affected by low MY, early to peak and 1-2 parity cows, low MY, mid to late and 1-2 parity cows, and low MY, early to peak and ≥ 3 parity cows (coefficients = −37.039, −25.353, −44.805 respectively, *P* < 0.05). When the cows were classified by MY before the heat season, the RT and MY of the high-MY group remained higher than the low-MY group until before the severe heat (THI ≥ 84 continued). However, when cows were classified by RT before the heat season, there was no difference in MY between the low-RT (< 485 min/d) and the high-RT group (> 485 min/d). In conclusions, MY is a factor affecting RT up to moderate heat but RT is not a sufficient condition affecting MY.

## 1 Introduction

Holstein dairy cows have low heat resistance because of their aria of origin (northern Germany and the Netherlands (Hurumura, 2012). Japan is hotter and more humid than these regions (Japan_Meteological_Agency, 2021), and the climate especially affects long-term heat stress to dairy cows in summer. The effects of heat stress include decreased production efficiency such as decreased appetite, increased number of diseases, and decreased milk production (Doukoshi, 2017; Sakatani, 2015). Thus, moderating heat stress to dairy cows in summer is necessary to prevent productivity decline in Japanese dairy farming.

Rumination is a physiological response that cows perform when they are at ease. And dairy cows usually ruminate 8 h/d (Adin et al., 2009). Rumination time (RT) varies depending on the health status of the cow and is an indicator of physical condition and surrounding environment. Previous studies reported that RT decreases as temperature humidity index (THI) increases, with high-producing cows and early lactation cows being more prone to this decrease (Moretti et al., 2017; Muschner-Siemens et al., 2020). The authors conducted a study at a dairy farm in Mie Prefecture, Japan, which was hotter and more humid in summer than the conditions of the above studies, and confirmed that herd rumination time was significantly reduced at a THI ≥ 84, which is severely hot according to Johnson’s. criteria (Johnson, 1962; Seto & Toba, 2022). The RT of heat-sensitive cows was lower than that of heat-tolerant cows before heat season. Thus, the authors hypothesized that the heat-sensitive cows could be estimated from the RT before the heat season, and conducted the verification. As a result, cows estimated to be heat-sensitive had lower RT during the hot season and a larger decrease in RT from just before the hot season than those estimated to be heat-tolerant (Seto & Toba, 2022). Based on these results, the authors proposed that the measurement of rumination time before the heat season could be a diagnostic method for heat-sensitive cows.

In the present study, cows from the same farm (no phase feeding) and the same period were researched, thus the environmental conditions of the herd were almost identical. However, since the analysis did not consider the conditions of the cows, such as parity, lactation at the beginning of the study, etc., it is not clear whether differences in these conditions make a difference in the tendency of the estimation of heat-sensitive cows before the heat period. Of course, it is not clear which of the above conditions has the greatest effect on heat sensitivity.

In this study, we compared the tendency of heat-sensitive cows by classifying the herds in the same farm according to the conditions of cows, and conducted a multivariate analysis using cow condition as the explanatory variable and RT before the heat season as the objective variable, in order to understand which condition is associated with RT.

## 2 Materials and Methods

### 2.1 Farm information

Data of cows were collected from the one dairy farm (Tsu, Mie, Japan). The milking capacity of this farm is 500 cows. The design of the barn is free stall and the milking method is by using milking robots (Astronaut A4, Lely, Netherlands). Cows fed partial mixed ration and could eat freely it. In addition, cows fed compound feed at their milking. To prevent the heat stress of cows, this farm used cheesecloth and curtains to block sunlight from entering the barn, fans to provide relay-type ventilation, a fine mist system to cool the air using vaporization heat. In addition, this farm used direct sprinkling showers to cool the cows during the daytime during periods of extreme heat.

### 2.2 Settings the research period

Weather data for Tsu, Mie, Japan from January to October 2019 was obtained from the Japan Meteorological Agency website (https://www.data.jma.go.jp/gmd/risk/obsdl/). THI were calculated from the maximum temperature and average humidity of this data. Heat season was defined from the start date when THI ≥ 72 continued for more than 7 d to the start date when THI ≤ 72 continued for more than 7 d. The research period was set from 7 d before the start of the heat season to the end of the heat season.

### 2.3 Data collection

RT (min/d) and milk yield (kg/d) during the research period, and parity and date of last calving at the date of data collection (October 23, 2019) were obtained from the herd management system (T4C management system, Lely, Netherlands) and the attached wearable sensor to measure RT (HR tag, SCR, Israel), and the milking robot. From the acquired data (total 533 cows), 253 cows that were in lactation at the beginning of the heat season and whose parity and days post calving were clear were used in this study. This data includes cows that were sick during the period. Note that data of RT, milk yield were included in the author’s present study (Seto & Toba, 2022).

### 2.4 Data preprocessing

Preprocessing of the collected data was performed for use in subsequent analyses. Files (each file contained RT, milk yield, parity, last calving date etc. of a cow) exported from the herd management system was converted from html format (looked xlsx format) to xlsx format. From converted files, the necessary data for this research period were extracted and each item was integrated into a single file as herd information. These preprocessing was performed using spreadsheet software (Microsoft Excel, Microsoft, USA), the programming language AppleScript (Apple, USA), the programming language python (Python Software Foundation, USA), and its libraries numpy (Harris et al., 2020) and pandas (McKinney, 2010). Sample computer scripts created for the preprocessing were uploaded on my repository of github (https://github.com/Takahiro-Seto/preprocessing_T4C_data).

### 2.5 Observation of each factor before heat season

From the pre-processed data, RT (min/d) of each cow during the 7 days before heat season were extracted and averages were calculated. From these average RT, samples were classified as low RT (< 485 min/d) or high RT (≥ 485 min/d) using the method of our present study (Seto & Toba, 2022). In the same way, milk yield (kg/d) of each cow during the 7 days before heat season were extracted and averages was calculated. These averages were then classified as low milk yield (< 45 kg/d) and high milk yield (≥ 45 kg/d) (Ibaraki_Prefecture, 2019). And the date of last calving was extracted and days post calving at the start of the hot season was calculated. From these days post calving, the lactation period at the start of the hot season of each cow were classified as early (equal or less than 60 d post calving), peak (more than 60 and equal or less than 90 d post calving), middle (more than 90 and equal or less than 210 d post calving) or late (more than 210 d post calving). The parity of each cow was classified as first, second or third or more (3≤).

From these classified data, a crosstabulation table was created with RT as the objective variable. In addition, histograms were created with milk yield, lactation period or parity as explanatory variables and rumination time as the objective variable.

### 2.6 Multivariate analysis

Using results of materials and methods 2.5, cows used in this study was divided into eight groups (group IDs and their element were described in Table 1). Group IDs were transformed into dummy variables, and multiple regression analysis using the dummy variables as explanatory variables (Hayashi’s quantification method-I) (Hayashi, 1954) was conducted with the average rumination time during the 7 days before the heat season as the objective variable. Multiple regression analysis was performed using a model with all dummy variables (Model 1) and a model with variable selection using Akaike’s information criterion (Model 2) (Akaike, 1973). These statistical analysis were performed by using the programming language R (R Core Team, 2020).

**Table 1.** Group ID and their elements for multivariate analysis

| Group ID | Average milk yield during 7 d before heat season (kg/d) | Lactation period at the start of heat season | Parity |
| --- | --- | --- | --- |
| Low_EP_12 | < 45 | early<sup>†</sup> or peak<sup>‡</sup> | first or second |
| Low_EP_3more | < 45 | early or peak | third or more |
| Low_ML_12 | < 45 | middle<sup>§</sup> or late<sup>¶</sup> | first or second |
| Low_ML_3more | < 45 | middle or late | third or more |
| High_EP_12 | ≥ 45 | early or peak | first or second |
| High_EP_3more | ≥ 45 | early or peak | third or more |
| High_ML_12 | ≥ 45 | middle or late | first or second |
| High_ML_3more | ≥ 45 | middle or late | third or more |

<sup>†</sup> equal or less than 60 d post calving
<sup>‡</sup> more than 60 and equal or less than 90 d post calving
<sup>§</sup> more than 90 and equal or less than 210 d post calving
<sup>¶</sup> more than 210 d post calving

### 2.7 Time-series observation of RT and milk yields during heat season

From the average milk yield during the 7 days before the heat season, the cows were classified into low (< 45 kg/d) and high (≥ 45 kg/d) milk yield groups. And the THI and RT or milk yield of the two groups during the heat season (from May 22 to October 13) were plotted.

In addition, from the average RT during the 7 days before the heat season, the cows were classified into low (< 485 kg/d) and high (≥ 485 kg/d) RT groups. And the THI and milk yield of the two groups during the heat season (from May 22 to October 13) were plotted.

## 3 Results

### 3.1 Research period

The research period was set from May 15 to October 13, 2019, with the start of the heat season on May 22.

### 3.2 Single element analysis by observing of crosstabulation table and histogram

The crosstabulation table with rumination time as the objective variable is shown in Table 2. When cows were classified according to average milk yield during the 7 d before the heat season, the percentage of cows with low average RT during the 7 d before the heat season was higher than 50% in the low milk yield group (54.1%). When the cows were classified according to lactation period at the beginning of the heat season (May 22), the percentage of cows with low average RT during the 7 days before the heat season was higher than 50% in all lactation period, in the order of early (58.1%), peak (57.1%), late (55.1%), and middle (50.3%). When the cows were classified according parity at the beginning of the heat season, the percentage of cows with low average RT during the 7 d before the heat season was higher than 50% in the first or ≥3 group.

**Table 2.** Crosstabulation table with rumination time as the objective variable

| Elements |  |  | Average RT<sup>†</sup> during the 7 days before heat season |  |  |
| --- | --- | --- | --- | --- | --- |
|  |  |  | Low RT<br>( < 485 min/d) | High RT<br>( ≥ 485 min/d) | Total |
| Average milk yield during the 7 d before heat season | Low milk yield (< 45 kg/d) | Headage | 98 | 83 | 181 |
|  |  | % | 54.1 | 45.9 | 100.0 |
|  | High milk yield (≥ 45 kg/d) | Headage | 36 | 36 | 72 |
|  |  | % | 50.0 | 50.0 | 100.0 |
| Lactation period at the start of heat season (5/22) | Early | Headage | 18 | 13 | 31 |
|  |  | % | 58.1 | 41.9 | 100.0 |
|  | Peak | Headage | 16 | 12 | 28 |
|  |  | % | 57.1 | 42.9 | 100.0 |
|  | Middle | Headage | 73 | 72 | 145 |
|  |  | % | 50.3 | 49.7 | 100 |
|  | Late | Headage | 27 | 22 | 49 |
|  |  | % | 55.1 | 44.9 | 100.0 |
| Parity at the start of heat season | 1 | Headage | 38 | 35 | 73 |
|  |  | % | 52.1 | 47.9 | 100.0 |
|  | 2 | Headage | 31 | 33 | 64 |
|  |  | % | 48.4 | 51.6 | 100.0 |
|  | ≥ 3 | Headage | 65 | 51 | 116 |
|  |  | % | 56.0 | 44.0 | 100.0 |

<sup>†</sup> Abbreviation: rumination time

Histograms of average RT during the 7 d before the heat season, with cows classified by average milk yield during the 7 d before the heat season, lactation period at the beginning of the heat season or parity, are shown in Figure 1. When the cows were classified by milk yield, the distribution of the high milk yield group showed two peaks (ranges 440-470 and 530-560 min/d), while the distribution of the low milk yield group showed a gentle distribution (peak was the range 440-470 min/d) (Figure 1a). When the cows were classified by lactation period, two peaks were clearly identified in the early (ranges 440-470 and 530-560 min/d) and peak (ranges 440-470 and 530-560 min/d). However, the middle stage distribution was more gentle (peak ranges 440-470 and 530-560 min/d) than the early and peak, and no clear peaks were identified in the late (Figure 1b). When the cows were classified by parity, no clear peaks were identified in the first parity. However, two peaks were identified in the second parity, and two peaks were clearer in the ≥3 parity.

**Figure 1.**
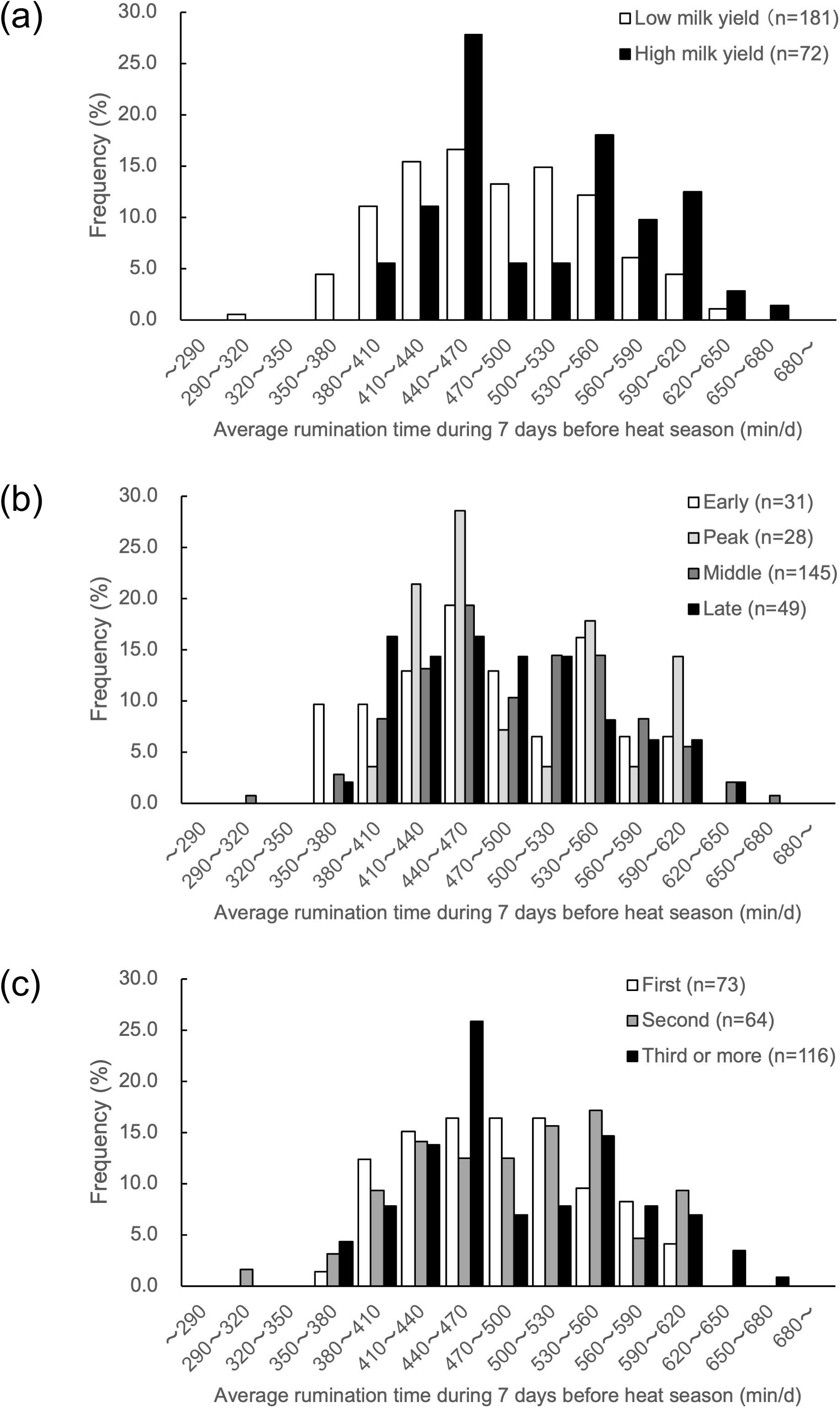
Histogram of Average rumination time (RT) during 7 d before heat season. (a) Comparison of distribution by milk yield. Low milk yield: Less than 45 kg/d of average milk yield during 7 d before heat season. High milk yield: 45 kg/d or more. (b) Comparison of distribution by lactation periods at the start of heat season. Early: 60 d or less. Peak: More than 60 and equal or less than 90 d. Middle: More than 90 and equal or less than 210 d. Late: More than 210 d. (c) Comparison of distribution by parity at the start of heat season.

### 3.3 Multivariate analysis

The cows in this study were classified eight groups based on Results 3.2 (Table 1). And multiple regression analysis was performed with each group as a dummy variable. Coefficients of Model 1 (used all dummy variables) and Model 2 (used 4 variables according to variable selection using AIC) are shown in Table 3. In Model 1, no dummy variable was found to have a significant effect on the multiple regression equation (*P* > 0.05). In model 2, dummy variable Low_EP_12, Low_ML_12, Low_EP_3more (meanings of these variable are shown in Table 1) were found have a significant negative effect on the multiple regression equation (*P* < 0.05).

**Table 3.** Coefficient of multiple regression analysis using dummy variable

|  | Dummy variable | Estimate | Standard error | t value | P |
| --- | --- | --- | --- | --- | --- |
| Model 1<br>(P of the model = 0.103) | (Intercept) | 500.565 | 10.816 | 46.280 | < 2<sup>-16</sup>* |
|  | Low_EP_12 | -31.266 | 17.170 | -1.821 | 0.070 |
|  | Low_EP_3more | -39.032 | 22.078 | -1.768 | 0.078 |
|  | Low_ML_12 | -19.580 | 12.857 | -0.523 | 0.129 |
|  | Low_ML_3more | -15.853 | 14.229 | -1.114 | 0.266 |
|  | High_EP_12 | 28.793 | 25.936 | 1.110 | 0.268 |
|  | High_EP_3more | -1.032 | 20.845 | -0.050 | 0.961 |
|  | High_ML_12 | 16.650 | 22.078 | 0.754 | 0.452 |
|  | High_ML_3more | NA<sup>†</sup> | NA | NA | NA |
| Model 2<br>(P of the model = 0.036) | (Intercept) | 506.338 | 7.837 | 64.608 | < 2<sup>-16</sup>* |
|  | Low_EP_12 | -37.039 | 15.437 | -2.399 | 0.0172** |
|  | Low_ML_12 | -25.353 | 10.464 | -2.423 | 0.0161** |
|  | Low_EP_3more | -44.805 | 20.735 | -2.161 | 0.0317** |
|  | Low_ML_3more | -21.626 | 12.102 | -1.787 | 0.0752 |

<sup>†</sup> Abbreviation: Not Available in R
\* p < 0.01
\*\* p < 0.05

### 3.4 Time-series observation of RT and milk yields during heat season

THI and RT during the heat season were shown in Figure 2a. Until mid-July, the RT of the high milk yield group (≥ 45 kg/d before heat season) remained higher than that of the low milk yield group (< 45 kg/d before heat season). However, from late-July, THI 84 or higher continued for more than two days, The RT of both was reduced less than 350 min/d and there was no difference between them. Thereafter, the RT of both fluctuated as THI increased or decreased, but the RT of the high milk yield group was not higher than that of the low milk yield group, as it was before mid-July.

**Figure 2.**
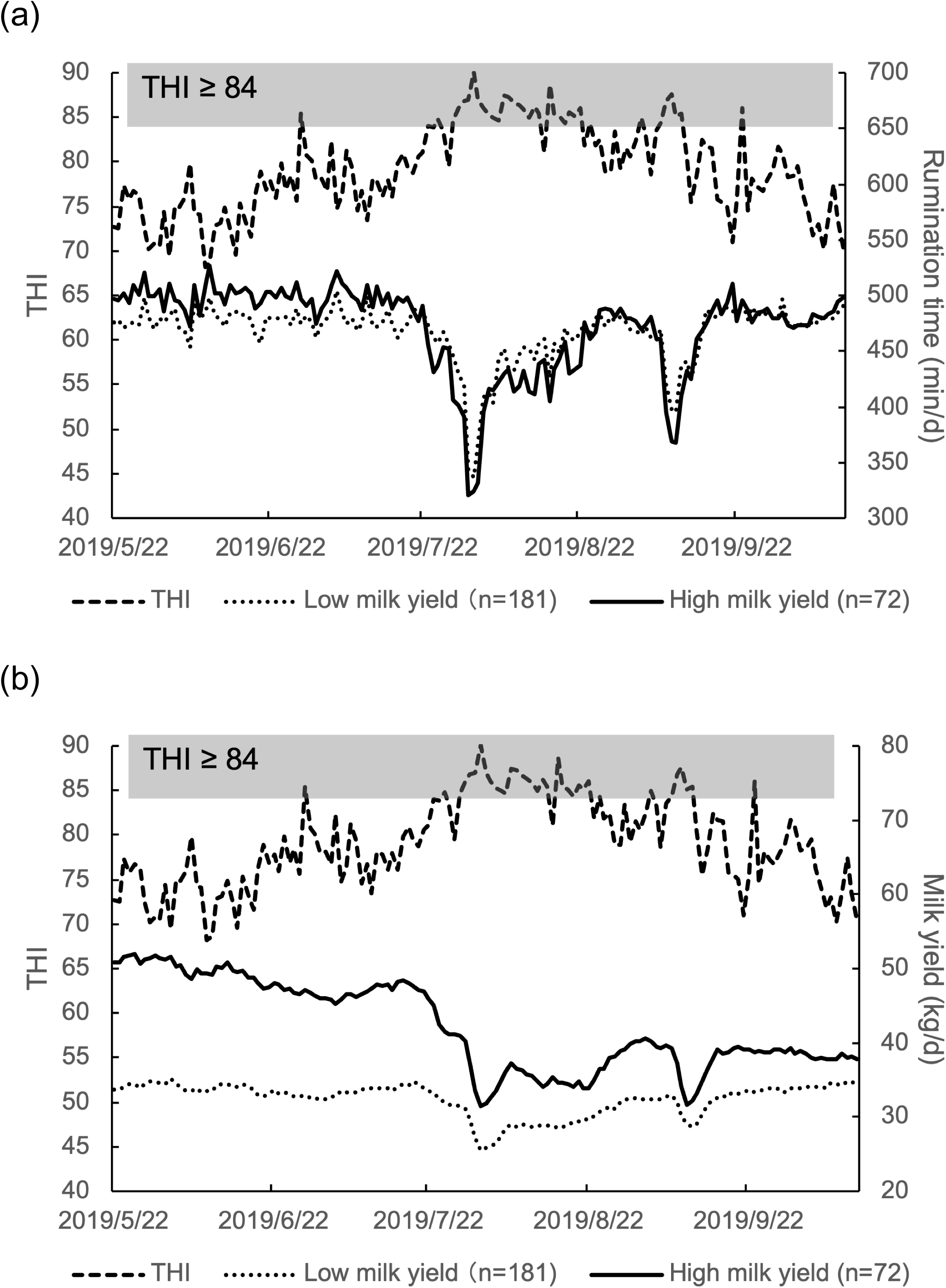
Graph of temperature humidity index (THI) and milk yield (a) or rumination time (b) in heat season. Low milk yield (n = 181): Less than 45 kg/d of average milk yield during 7 d before heat season. High milk yield (n = 72): 45 kg/d or more.

THI and milk yield during the heat season were shown in Figure 2b and 3. The milk yield of the high milk yield group remained higher than that of the low milk yield group during the all-heat season. However, from late-July, the milk yield of both was reduced significantly and fluctuated as THI increased or decreased. From September, THI 84 or higher did not continue for more than two days, the milk yield in the low milk yield group recovered to the same level as before mid-July. However, the milk yield in the high milk yield group did not recover (Figure 2b). On the other hand, the milk yield of high (≥ 485 min/d before heat season) and low (< 485 min/d before heat season) RT groups remained similar during the all-heat season (Figure 3).

**Figure 3.**
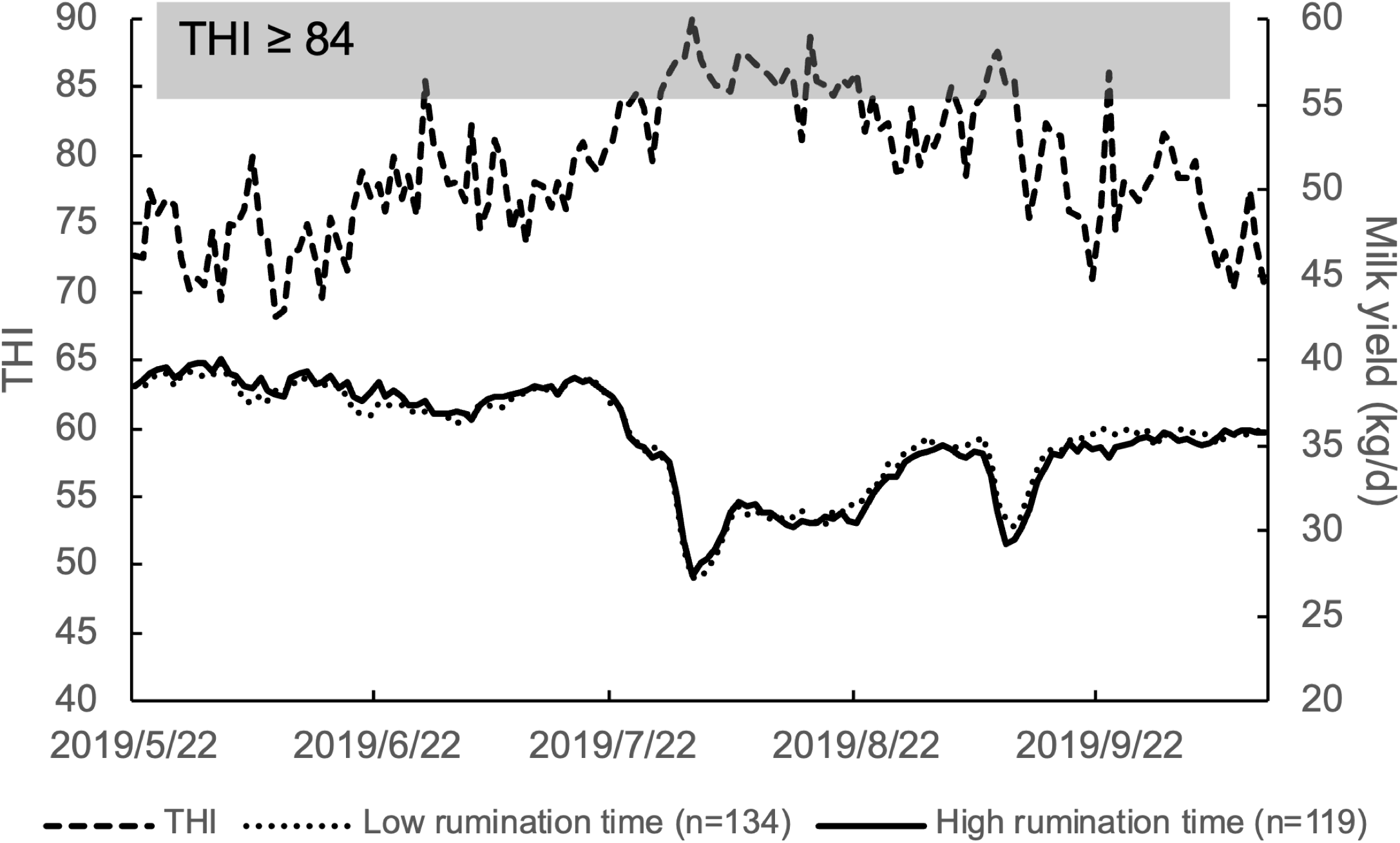
Graph of temperature humidity index (THI) and milk yield in heat season. Low rumination time: Less than 485 min/d of average rumination time during 7 d before heat season. High rumination time: 485 min/d or more.

## 4 Discussion

The summer climate in Japan is very hot and humid even compared to tropical regions, with persistent THI above 84, which is defined severe heat by Johnson’s criteria (Johnson, 1962). Therefore, it is very important in dairy management in Japan to avoid heat stress in summer. It is to be desired to detect heat-sensitive cows at an early stage because it naturally takes time to implement heat control measures. The author’s previous study proposed a method to estimate heat-sensitive cows based on RT before the heat season (Seto & Toba, 2022). However, in this study, the analysis was conducted without considering cow attributes such as milk yield, lactation period, and parity. Thus, in this study, cow attributes influenced RT before the heat season were analyzed.

Analysis by observation of crosstabulation tables showed that herds with low milk yield before the heat season, early, peak, or late lactation at the start of the heat season, and first or ≥3 parity at the start of the heat period tended to have low RT before the heat season. Analysis by observation of histograms also confirmed differences in distribution between low and high milk yield, early or peak lactation and mid or late lactation period, and first or second and ≥3 parity. In general, milk yield is higher in early or peak lactation than in mid or late lactation, and the milk yield increases up to third parity (Lee & Kim, 2006; Maltz et al., 1991). Thus, these three factors are causally related and it is difficult to explore the causal relationship with ruminant time by univariate analysis. On the other hand, the similarity between the results of the crosstabulation table and the histogram suggests that at least for milk yield and milk period, it is possible to divide within attributes into two categories: low milk yield and high milk yield, early and peak period and mid and late period. In terms of parity, the distribution of second calving was intermediate between the distribution of first and ≥3 parity, while the percentage of cows with low RT before heat season was the lowest. During the transition period, first parity cows are known to have different feeding behavior compared to high parity cows (Neave et al., 2017). However, we did not understand why there were fewer low RT cows in the second parity. In addition to feeding behavior, the movement behavior of dairy cows changes as the number of parity increases (Vázquez Diosdado et al., 2018). Considering these previous reports and the results of histograms, cows were classified into first or second and ≥3 parity. As described above, the cows were divided into eight groups by multiplying the binary classification of milk yield, lactation, and calving age, and multiple regression analysis using dummy variables was conducted.

The results of multiple regression analysis suggested that low milk yield affected RT in the whole lactation periods of first and second parity, and in the early and peak periods for ≥3 parity. The p-value for the middle and late stages of the hot season was close to the statistically significant p-value (*P* = 0.0752). These results suggest that low milk yield may be due to low rumination in the pre-heat season.

Based on the results of the multiple regression analysis, we classified the cows by milk yield before the heat season and observed the changes in RT and milk yield during the heat season. As THI increased, ruminating time and milk yield decreased, and as THI decreased, ruminating time and milk yield increased, until just before the latter half of July, when THI was 84 or higher continued, which is severe heat. In both cases, the high milk yield group before heat season remained higher than the low milk yield group. However, after the latter half of July, as THI increased, ruminating time and milk yield of the high milk yield group both decreased to the same level as that of the low milk yield group. And even when THI decreased due to seasonal changes, RT and milk yield did not recover to the same level as before the latter half of July, especially RT remained at the same level as that of the low milk yield group. On the other hand, RT and milk yield of the low milk yield group recovered to the same level as before the latter half of July when THI came down even after being affected by severe heat. These results suggest that severe heat above THI 84 has an irreversible effect on high milk yield cows, especially reflected in RT. The high temperature itself irreversibly interferes with the ability of mammary gland cells to synthesize milk (Kobayashi, Kawahara & Isobe, 2020; Sun et al., 2021), and high milk yielding cows also generate more heat due to metabolism (National Agrifulture and Food Research Organization, 2017). This suggests that severe heat irreversibly inhibits the ability of mammary gland cells to synthesize milk, especially in high producing cows. Another possible explanation is a decrease in milk production due to changes over time in the lactation period, since milk production is higher in early and peak lactation compared to mid and late lactation. Further analysis of these issues is needed. On the other hand, we did not understand why the ruminating time of the high milk yield group remained at the same level as that of the low milk yield group even when heat improved. According to the member of the research farm, the time when heat damage to cows was completely cured was in January. Therefore, it may be necessary to observe cows longer than the research period in this study.

In order to verify whether low RT before the heat period affects low milk yield, we classified cows into RT before heat season and observed milk yield trends during the heat season. This suggested that low RT did not affect low milk yield. When discussed together with the results of the multiple regression analysis, the following logical equation was obtained in moderate heat or less.

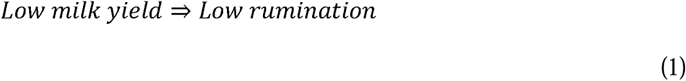

The hypothesis of the mechanism of the above is described below. In high lactation dairy cows, postpartum energy requirements are more than three times higher than maintenance (National Agrifulture and Food Research Organization, 2017). This indicates that most of the energy ingested by lactating cows is used for milk production. Therefore, the effect of milk yield on energy requirements is greater than other factors such as age and body weight. And the energy requirements of low-milking cows are less than those of high-milking cows. The factors that determine milk production of cows, in the same farm and at the same time, are the lactation performance of the cows themselves, in addition to the parity, lactation period and nutrition of the food consumed. Factors affecting lactation performance include rumen function, liver function, and milk synthesis in the mammary gland (Oba, 2019). However, if the rumen function affects lactation performance in this study, it is difficult to discuss that rumination does not influence digestion in the rumen. On the other hand, milk synthesis capacity in the mammary gland is directly related to genetic factors, lactation, and nutritional status, but not to rumen function (via liver function and nutritional status). Therefore, it is likely that energy requirements are controlled by milk synthesis capacity in the mammary gland, which in turn controls rumen function and rumination time.

In conclusion, at least below moderate heat, low milk production is a factor for low rumination, but low rumination is not a factor for low milk production, and severe heat has irreversible effects on rumination time and milk production, especially for high milk producing cows. In terms of feed efficiency, high milk yielding cows are more desirable. However, RT did not recover as same as milk yield in high milk yielding cow after severe heat. This may cause malnutrition in cows due to slow digestion of feed. While farm-gate milk price of japan is higher than other countries (CLAL, 2022), it also has a low feed self-sufficiency rate (Japan Ministry of Agriculture, Forestry and Fisheries, 2014). Due to the logistics disruption from COVID-19 (World Trade Organization, 2022) and climate change (IPCC, 2021), it is becoming difficult to procure purchased feed. And even about self-sufficient feed only, what can be grown varies by region, making it difficult to supply sufficient nutrition to cows. Feeding such as bypass amino acids can be considered to provide the nutrition that is lacking in feed, but considering the cost effectiveness, it is not always possible to provide the best nutrition for cattle. Furthermore, how much heat countermeasures are taken, it is impossible to escape from severe heat itself. Under these circumstances, I think it is important to consider what cows are better for each farm in future dairy management.

## Acknowledgements

We thank the Yamagishism Seikatsu Toyosato Jikkenchi Agricultural Cooperative Corporation (Mie, Japan) for cooperating as a research farm.

## Ethical considerations

This study was conducted in accordance with the guidelines set of Shizuoka Professional University Junior College of Agriculture.

Because this research does not involve human subjects, this study has not been reviewed by the Shizuoka Professional University Junior College of Agriculture Ethics Code (For research involving direct human subjects, conformed to the rules of the Declaration of Helsinki). However, in compliance with this code, we are handling the data obtained from the research so that it will not be leaked.

